# ProSelfLC: Progressive Self Label Correction Towards A Low-Temperature Entropy State

**DOI:** 10.1101/2022.07.01.498447

**Authors:** Xinshao Wang, Yang Hua, Elyor Kodirov, Sankha Subhra Mukherjee, David A. Clifton, Neil M. Robertson

## Abstract

To train robust deep neural networks (DNNs), we systematically study several target modification approaches, which include output regularisation, self and non-self label correction (LC). Three key issues are discovered: (1) Self LC is the most appealing as it exploits its own knowledge and requires no extra models. However, how to automatically decide the trust degree of a learner as training goes is not well answered in the literature. (2) Some methods penalise while the others reward low-entropy predictions, prompting us to ask which one is better. (3) Using the standard training setting, a trained network is of low confidence when severe noise exists, making it hard to leverage its high-entropy self knowledge.

To resolve the issue (1), taking two well-accepted propositions–deep neural networks learn meaningful patterns before fitting noise and minimum entropy regularisation principle–we propose a novel end-to-end method named ProSelfLC, which is designed according to learning time and entropy. Specifically, given a data point, we progressively increase trust in its predicted label distribution versus its annotated one if a model has been trained for enough time and the prediction is of low entropy (high confidence). For the issue (2), according to ProSelfLC, we empirically prove that it is better to redefine a meaningful low-entropy status and optimise the learner toward it. This serves as a defence of entropy minimisation. To address the issue (3), we decrease the entropy of self knowledge using a low temperature before exploiting it to correct labels, so that the revised labels redefine a low-entropy target state.

We demonstrate the effectiveness of ProSelfLC through extensive experiments in both clean and noisy settings, and on both image and protein datasets. Furthermore, our source code is available at https://github.com/XinshaoAmosWang/ProSelfLC-AT.

## 1 Introduction

THERE exist many target (label) modification approaches. They can be roughly divided into two groups: (1) Output regularisation (OR), which is proposed to penalise overconfident predictions for regularising deep neural networks. It includes label smoothing (LS) [55, 72] and confidence penalty (CP) [59]; (2) Label correction (LC). On the one hand, LC regularises neural networks by adding the similarity structure information over training classes into one-hot label distributions so that the learning targets become *structured and soft*. On the other hand, it can *correct the semantic classes* of noisy label distributions. LC can be further divided into two subgroups: Non-self LC and Self LC. The former requires extra learners, while the latter relies on the model itself. A typical approach of Non-self LC is knowledge distillation (KD), which exploits the predictions of other model(s), usually termed teacher(s) [30]. Self LC methods include Pseudo-Label [40], bootstrapping (Boot-soft and Boot-hard) [62], Joint Optimisation (Joint-soft and Joint-hard) [73], and Tf-KD_*self*_ [97]. According to an overview in Fig. 1 (detailed derivation is in Section 3 and Table 1), *in label modification, the output target of a data point is defined by combining a one-hot label distribution and its corresponding prediction or a predefined label distribution*.

**TABLE 1:**
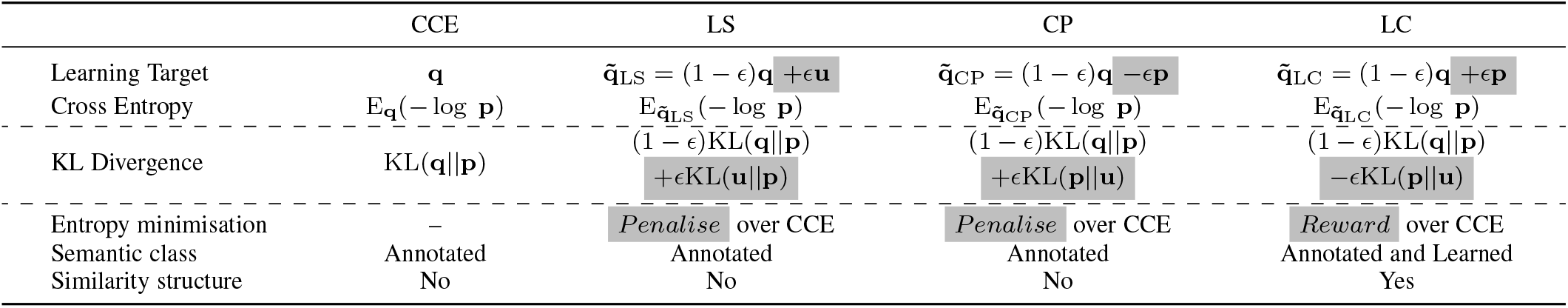
Summary of CCE, LS, CP and LC.

**Fig. 1:**
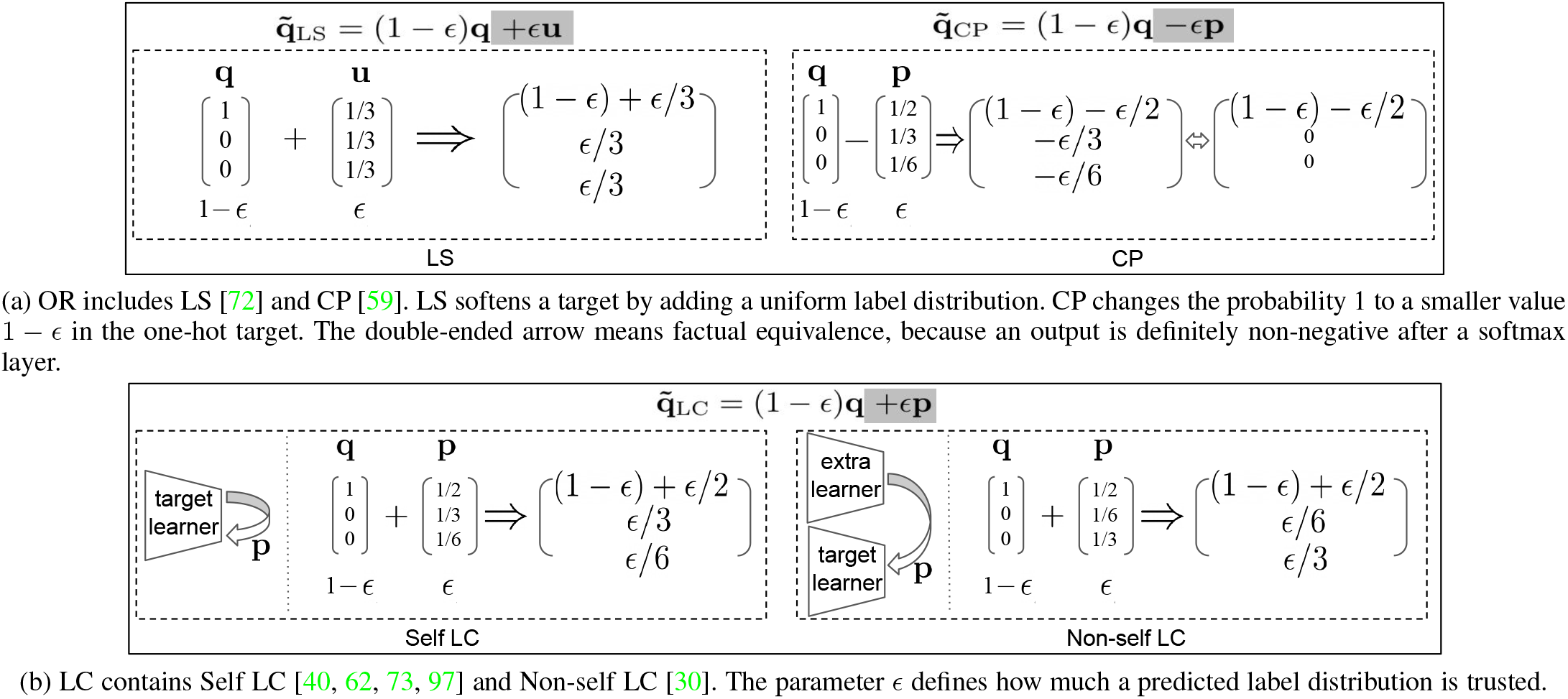
Target modification includes OR (LS and CP), and LC (Self LC and Non-self LC). Assume there are three training classes. **q** is the one-hot target. **u** is a uniform label distribution. **p** denotes a predicted label distribution. The target combination parameter is *ϵ* ∈ [0, 1].

Firstly, we present the drawbacks of existing approaches: (1) OR methods naively penalise confident outputs without leveraging easily accessible knowledge from other learners or itself (Fig. 1a); (2) Non-self LC relies on accurate auxiliary models to generate predictions (Fig. 1b). (3) Self LC is the most appealing because it exploits its own knowledge and requires no extra learners. However, there is a core question that is not well answered:

### In Self LC, how much should we trust a learner to leverage its knowledge?

As shown in Fig. 1b, in Self LC, for a data point, we have two labels: a predefined one-hot **q** and a predicted structured **p** (i.e., self knowledge). Its learning target is (1 − *ϵ*)**q** + *ϵ* **p**, i.e., a trade-off between **q** and **p**, where *ϵ* defines the trust score of a learner. In existing methods, *ϵ* is fixed without considering that a model’s knowledge grows as the training progresses. For example, in bootstrapping, *ϵ* is fixed throughout the training process. Joint Optimisation stage-wisely trains a model. It fully trusts predicted labels and uses them to replace old ones when a stage ends, i.e., *ϵ* = 1. Tf-KD_*self*_ trains a model by two stages: *ϵ* = 0 in the first one while *ϵ* is tuned for the second one. Note that **p** is generated by a preceding-stage model in stage-wise training, which requires significant human intervention and is time-consuming in practice.

To improve Self LC, we propose a novel method named Progressive Self Label Correction (ProSelfLC), which is end-to-end trainable and needs negligible extra cost. Most importantly, ProSelfLC modifies the target progressively and adaptively as training goes. *Two design principles of ProSelfLC are*: (1) When a model learns from scratch, human annotations are more reliable than its own predictions in the early phase, during which the model is learning simple meaningful patterns before fitting noise, even when severe label noise exists in human annotations [4]. (2) As a learner attains confident knowledge as time progresses, we leverage it to revise annotated labels. This is surrounded by minimum entropy regularisation, which is widely evaluated in unsupervised and semi-supervised scenarios [20, 21].

Secondly, note that OR methods penalise low entropy while LC rewards it, intuitively leading to the second vital question:

### Should we penalise a low-entropy status or reward it?

Entropy minimisation is the most widely used principle in machine learning [20, 21, 27, 39, 66]. In standard classification, minimising categorical cross entropy (CCE) optimises a model towards a low-entropy status defined by human annotations, which contain noise in very large-scale machine learning. As a result, confidence penalty becomes popular for reducing noisy fitting. In contrast, we prove that it is better to reward a meaningful low-entropy status redefined by our ProSelfLC. Therefore, our work offers a defence of entropy minimisation against the recent confidence penalty practice [16, 55, 59, 72].

Thirdly, we highlight a common phenomenon which hinders a model from confident learning towards a low-entropy target state. By reporting the confidence metrics in Fig. 2 and Table 9, we display this phenomenon, i.e., *a deep model learns much less confidently when the training data contains more severe noise*. Inspired by this, to redefine a corrected low-entropy target state and to reward a model’s confident learning, we develop an Annealed Temperature (AT) as a plug-in module to reduce the entropy of self knowledge **p**. Empirically (see Table 9), for the Self LC methods including Boot-soft and ProSelfLC, with an AT plugged in, we are able to exploit the low-temperature (i.e., low-entropy) self knowledge. Consequently, the models learn more confidently and generalise better.

**Fig. 2:**
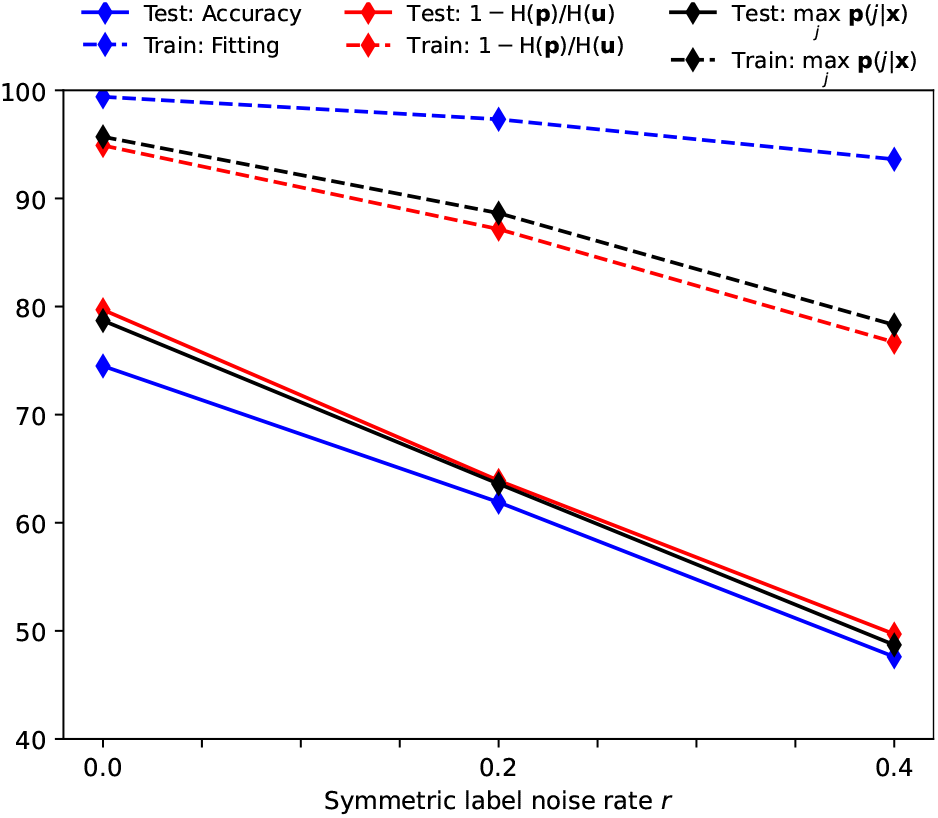
The changes of fitting train set, generalising on test set, and confidence metrics on both sets along with the symmetric label noise rate *r*. Two confidence metrics are shown using the red and black lines. The vertical axis shows the percentage of a metric’s result. We train ResNet18 on CIFAR-100 using the standard CCE. The final model is used for analysis when the training stops. Our finding: as *r* increases, though the fitting of train set decreases little [99], *the confidences of both sets drop significantly*.

We summarise our main contributions as follows:

- We provide a theoretical study on popular target modification methods through entropy and KL divergence [37]. Accordingly, we reveal their drawbacks and propose ProSelfLC as a solution. ProSelfLC can: (1) enhance the similarity structure information over training classes; (2) correct the semantic classes of noisy label distributions. ProSelfLC is the first method to progressively and adaptively trust a low-temperature self knowledge.
- We present a new finding which is complementary to [99]: *Deep models are significantly less confident when higher label noise exists*. Correspondingly, we propose to decrease the entropy of self knowledge using an AT and learn towards a revised low-temperature entropy state.
- Our extensive experiments: (1) defend the entropy minimisation principle; (2) demonstrate ProSelfLC’s effectiveness in clean and noisy settings of two very diverse data domains, i.e., image and protein datasets. This demonstrates the general applicability of our method.

## 2 Related Work

### Label noise and semi-supervised learning

We test target modification approaches in the setting of label noise because it is generic and connected with semi-supervised learning, where only a subset of training examples are annotated, leading to *missing labels*. Then the key to semi-supervised training is to reliably fill them. *When these missing labels are incorrectly filled, the challenge of semi-supervised learning changes to noisy labels*. For a further comparison, in semi-supervised learning, the annotated set is clean and reliable, because the label noise only exists in the unannotated set. While in our experimental setting, we are not given information on whether an example is trusted or not, thus being even more challenging. We summarise existing common approaches for solving label noise: (1) Loss correction, in which we are given or we need to estimate a noise-transition matrix, which defines the distribution of noise labels [13, 19, 24, 33, 45, 46, 50, 58, 71, 74, 85–87, 92, 103, 109]. A noise-transition matrix is difficult and complex to estimate in practice; (2) Exploiting an auxiliary trusted training set to differentiate examples [29, 42, 75]. This requires extra annotation cost; (3) Co-training strategies, which train two or more learners [25, 32, 43, 54, 60, 80, 96] and exploit their ‘disagreement’ information to differentiate data points; (4) Label engineering methods [40, 43, 62, 68, 73, 92, 95, 105], which relate to our focus in this work. Their objective is to annotate unlabelled samples or correct noisy labels; (5) Sample selection methods aim to drop corrupted examples [14, 81, 84, 93]; (6) Example weighting approaches assign higher weights to more reliable labels [6, 10, 23, 32, 38, 63, 67, 77]; (7) Robust loss functions include [48, 52, 76, 79, 106]. Most of them are compared in the experimental section.

### LC and knowledge distillation (KD)

[9, 30]. Mathematically, we derive that some KD methods also modify labels. We use the term label correction instead of KD for two reasons: (1) label correction is more descriptive; (2) the scope of KD is not limited to label modification. For example, multiple networks are trained for KD [18]. When two models are trained, the consistency between their predictions of a data point is promoted in [5, 104], while the distance between their feature maps is reduced in [65]. Regarding self KD, two examples of the same class are constrained to have consistent output distributions [91, 98]. In another self KD [101], the deepest classifier provides knowledge for shallower classifiers. In a recent self KD method [97], Tf-KD_*self*_ applies two-stage training. In the second stage, a model is trained by exploiting its knowledge learned in the first stage. Our focus is to improve the end-to-end self LC. First, self KD methods [91, 98, 101] maximise the consistency of intraclass images’ predictions or the consistency of different classifiers. In our view, they do not modify labels, thus being less relevant for comparison. Second, the two-stage self KD method [97] can be an add-on (i.e., an enhancement plugin) other than a competitor. E.g., in real-world practice, the first stage can be ProSelfLC instead of CCE with early stopping. Finally, we acknowledge that exploiting ProSelfLC to improve non-self KD and stage-wise approaches is an important area for future work, e.g., a better teacher model can be trained using ProSelfLC.

## 3 Mathematical Analysis and Theory

Let 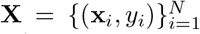 represent *N* training examples, where (**x**_*i*_, *y*_*i*_) denotes *i* −th sample with input **x**_*i*_ ∈ ℝ^*D*^ and label *y*_*i*_ ∈ { 1, 2, …, *C* }. *C* is the number of classes. A deep neural network *z* consists of an embedding network *f* (·) : ℝ^*D*^ →ℝ^*K*^ and a linear classifier *g*(·) : ℝ^*K*^ →ℝ^*C*^, i.e., **z**_*i*_ = *z*(**x**_*i*_) = *g*(*f* (**x**_*i*_)) : ℝ^*D*^ →ℝ^*C*^. For the brevity of analysis, we take a data point and omit its subscript so that it is denoted by (**x**, *y*). The linear classifier is usually the last fully-connected layer. Its output is named logit vector **z** ∈ ℝ^*C*^. We produce its classification probabilities **p** by normalising the logits using a softmax function:

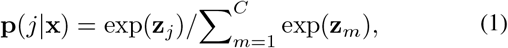

where **p**(*j* |**x**) is the probability of **x** belonging to class *j*. Its corresponding ground-truth is usually denoted by a one-hot representation **q**: **q**(*j*|**x**) = 1 if *j* = *y*, **q**(*j*|**x**) = 0 otherwise.

### 3.1 Semantic class and similarity structure in p

A probability vector **p** ∈ ℝ^*C*^ can also be interpreted as an instance-to-classes similarity vector, i.e., **p**(*j* | **x**) measures how much a data point **x** is similar with (analogously, likely to be) *j*-th class. Consequently, **p** should not be exactly one-hot, and is proposed to be corrected at training, so that it can define a more informative and structured learning target. For better clarity, we first present two definitions:

#### Definition 1

(*Semantic Class*). Given a target label distribution 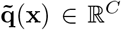, the semantic class is defined by 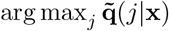, i.e., the class whose probability is the largest.

#### Definition 2

(*Similarity Structure*). In 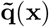, **x** has *C* probabilities of being predicted to *C* classes. The similarity structure of **x** versus *C* classes is defined by these probabilities and their differences.

### 3.2 Revisit of CCE, LS, CP and LC

#### Standard CCE

For any input (**x**, *y*), the minimisation objective of standard CCE is:

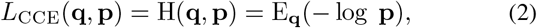

where H(·, ·) represents the cross entropy. E_**q**_(− log **p**) denotes the expectation of negative log-likelihood, and **q** serves as the probability mass function.

#### Label smoothing

In LS [30, 72], we soften one-hot targets by adding a uniform distribution: 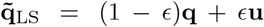, **u** ∈ ℝ^*C*^, and 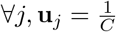. Consequently:

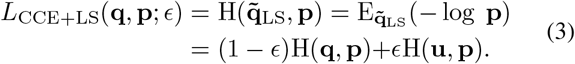

#### Confidence penalty

CP [59] penalises highly confident predictions:

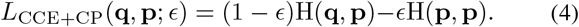

#### Label correction

As illustrated in Fig. 1, LC is a family of algorithms, where a one-hot label distribution is modified to a convex combination of itself and a predicted distribution:

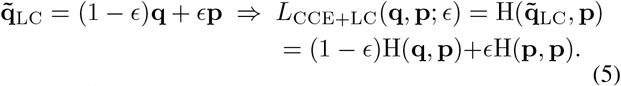

We remark: (1) **p** provides meaningful information about an example’s relative probabilities of being different training classes; (2) If *ϵ* is large, and **p** is confident in predicting a different class, i.e., arg max_*j*_ **p**(*j*|**x**) ≠ arg max_*j*_ **q**(*j*|**x**), 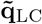 defines a different semantic class from **q**.

### 3.3 Theory

#### Proposition 1.

*LS, CP and LC modify the learning targets of standard CCE*.

*Proof. L*_CCE+CP_(**q, p**; *ϵ*) = (1 − *ϵ*)H(**q, p**) − *ϵ* H(**p, p**) = E_(**1**_ − _*ϵ*)**q**_ − _*ϵ* **p**_(− log **p**). Therefore,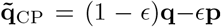. Addi-tionally, 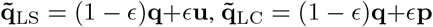. □

#### Proposition 2.

*Some KD methods, which aim to minimise the KL divergence between predictions of a teacher and a student, belong to the family of label correction*.

*Proof*. In general, a loss function of such methods can be defined to be *L*_KD_(**q, p**_*t*_, **p**) = (1 − *ϵ*)H(**q, p**) + *ϵ* KL(**p**_*t*_ || **p**) [97]. KL(·|| ·) denotes the KL divergence. As KL(**p**_*t*_ || **p**) = H(**p**_*t*_, **p**) − H(**p**_*t*_, **p**_*t*_), **p**_*t*_ is from a teacher and fixed when training a student. We can omit H(**p**_*t*_, **p**_*t*_):

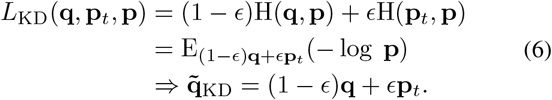

Consistent with LC in Eq (5), *L*_KD_(**q, p**_*t*_, **p**) revises a label using **p**_*t*_. □

#### Proposition 3.

*Compared with CCE, LS and CP penalise entropy minimisation while LC reward it*.

#### Proposition 4.

*In CCE, LS and CP, a data point* **x** *has the same semantic class. In addition*, **x** *has an identical probability of belonging to other classes except for its semantic class*.

The proofs of propositions 3 and 4 are presented in the Appendix A. Only LC exploits informative information and has the ability to correct labels, while LS and CP only relax the hard targets. We summarise CCE, LS, CP and LC in Table 1. Constant terms are ignored for concision.

## 4 ProSelfLC: Progressive and Adaptive Label Correction

In standard CCE, a semantic class is considered while the similarity structure is ignored. It is mainly due to the difficulty of annotating the similarity structure for every data point, especially when *C* is large [90]. Fortunately, recent progress demonstrates that there are some effective approaches to define the similarity structure of data points without annotation: (1) In KD, an auxiliary teacher model can provide a student model the similarity structure information [30, 55]; (2) In Self LC, e.g., Boot-soft, a model helps itself by exploiting the knowledge it has learned so far. We focus on studying the end-to-end Self LC to further comprehend and improve it in this work.

In Self LC, *ϵ* indicates how much a predicted label distribution is trusted. In ProSelfLC, we propose to set it adaptively according to the learning time *t* and confidence of **p**. For any **x**, we summarise its equations of loss *L*, label and trust below.

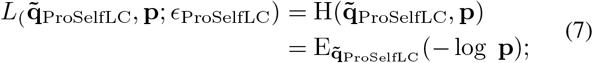

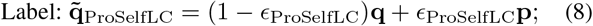

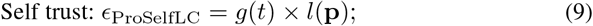

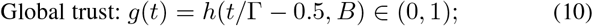

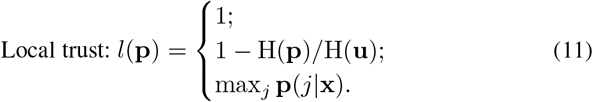

*t* and Γ are the iteration counter and the number of total iterations, respectively. *h*(*η, B*) = 1*/*(1 + exp(−*η* × *B*)). Here, *η* = *t/*Γ− 0.5. *B*, Γ are task-dependent and can be chosen according to a validation set in practice. In Eq. (11), we display three options to compute the local trust *l*(**p**).

### 4.1 Progressive and adaptive self trust by *ϵ* _ProSelfLC_

**Global trust** *g*(*t*) denotes how much we trust a learner. It is independent of data points, thus being global. *g*(*t*) grows as *t* rises. *B* adjusts the exponentiation’s base and growth speed of *g*(*t*). Theoretically and practically, *g*(*t*) could be many other formats, e.g., a stepwise growth, a linear increase, or a polynomial rise, etc. In the scope of this research, we only consider the exponential growth and leave other increase options for future work.

**Local trust** *l*(**p**) indicates how much we trust an output distribution **p**. If *l*(**p**) = 1, all predictions are treated equally. When *l*(**p**) = 1 − H(**p**)*/*H(**u**) or max_*j*_ **p**(*j* | **x**), a more confident prediction has a higher trust. 1 − H(**p**)*/*H(**u**) and max_*j*_ **p**(*j* | **x**) are different confidence metrics [22].

We will empirically discuss *g*(*t*) using different *B* and three *l*(**p**) options in the Section 6.5, where confidence-based 1 − H(**p**)*/*H(**u**) and max_*j*_ **p**(*j*|**x**) work better.

### 4.2 Design reasons

The design of *g*(*t*) is inspired by human learning. In the earlier learning phase, i.e., *t <* Γ*/*2, *g*(*t*) *<* 0.5 ⇒ *ϵ* _ProSelfLC_ *<* 0.5, ∀**p**, so that the predefined supervision dominates and ProSelfLC only modifies the similarity structure a bit. When a learner has not seen the training data for enough time at the earlier stage, its knowledge is less reliable and wrongly confident predictions can be produced. Our design assuages the bad impact of such undesirable cases. When it comes to the later training phase, i.e., *t >* Γ*/*2, we have *g*(*t*) *>* 0.5 as it has been trained for more than half of entire learning time.

Regarding *l*(**p**), it affects the later learning phase a lot. If **p** is less confident (i.e., of higher entropy), *l*(**p**) is smaller. Hence, *ϵ* _ProSelfLC_ is lower and we trust **p** less. If **p** is highly confident, *ϵ* _ProSelfLC_ is larger so we trust its confident knowledge.

### 4.3 Case analysis

Due to the potential memorisation in the earlier phase (though less likely to be severe), we may get undesired confidently wrong predictions for noisy labels, but their trust scores are small as *g*(*t*) is small. We conduct the case analysis of ProSelfLC in Table 2 and summarise its core tactics as follows:

**TABLE 2:** The values of *ϵ* _ProSelfLC_ = *g*(*t*) × *l*(**p**) under different cases. We use concrete values for concise interpretation. We bold the special case when the semantic class is changed. Consistency is defined by whether **p** and **q** share the semantic class or not.

| | | $l(\mathbf{p}) = 0.1$<br>(non-confident) | $l(\mathbf{p}) = 0.9$<br>(confidently consistent) | $l(\mathbf{p}) = 0.9$<br>(confidently inconsistent) |
| --- | --- | --- | --- | --- |
| Earlier phase | $g(t) = 0.1$ | 0.01 | 0.09 | 0.09 |
| Later phase | $g(t) = 0.9$ | 0.09 | 0.81 | **0.81** |

(1) Correct the similarity structure for every data point in all cases, thanks to exploiting the self knowledge of a learner, i.e., **p**.

(2) Revise the semantic class when *t* is large enough and **p** is confidently inconsistent. As highlighted in Table 2, when two conditions are met, we have *ϵ* _ProSelfLC_ *>* 0.5 and arg max_*j*_ **p**(*j* | **x**) ≠ arg max_*j*_ **q**(*j* | **x**), then **p** redefines the semantic class. For example, if **p** = [0.95, 0.01, 0.04], 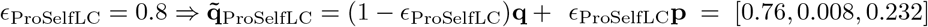. Note that ProSelfLC also becomes robust against lengthy exposure to the noisy data, as empirically demonstrated in Fig. 3 and Table 4.

**Fig. 3:**
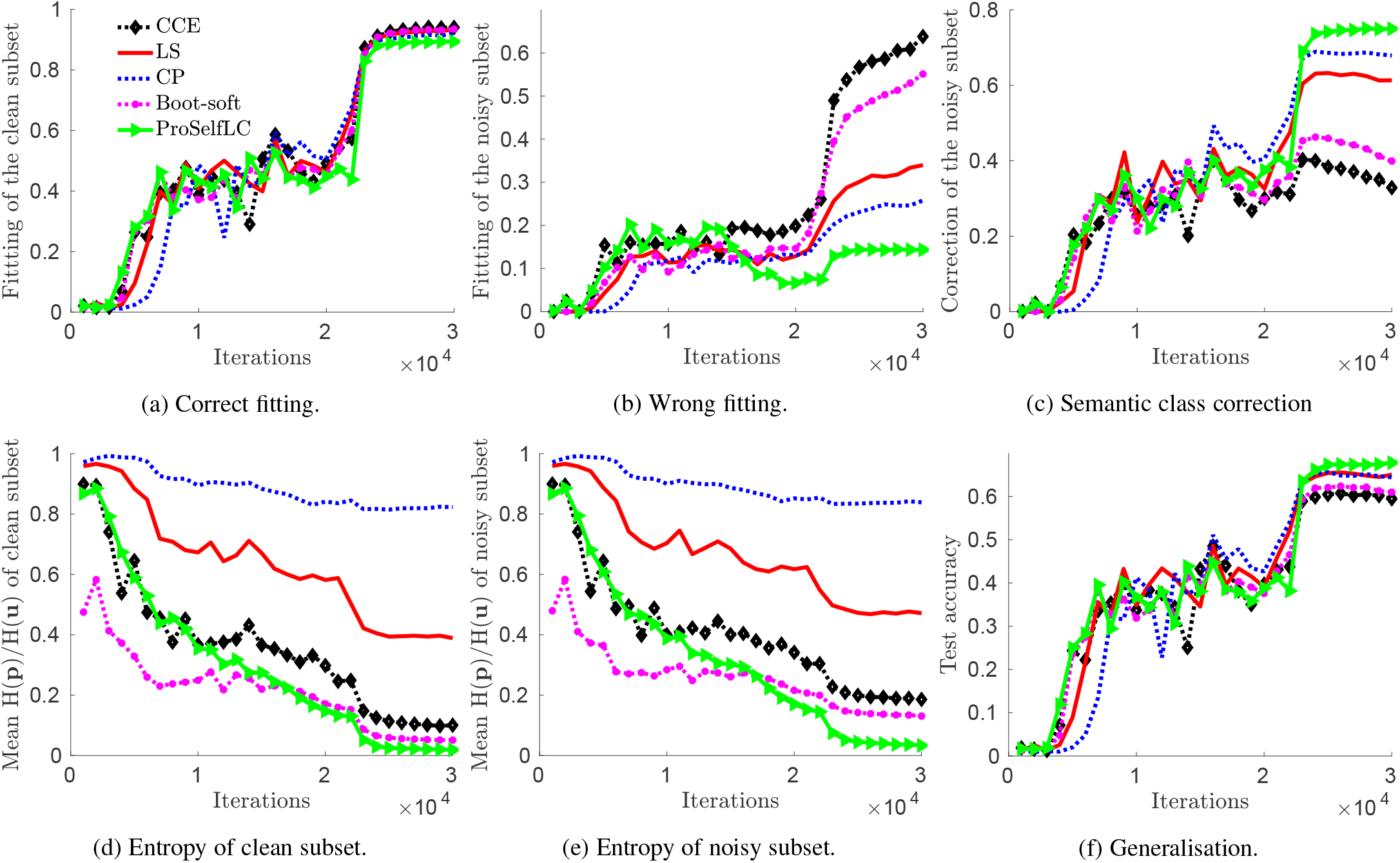
Dynamic learning statistics on CIFAR-100 with 40% asymmetric label noise. *At training, a learner is NOT GIVEN whether a label is trusted or not*. Therefore, the target modification methods can be treated as unsupervised. We store all intermediate models and plot their results of six metrics. Vertical axes and subcaptions describe evaluation metrics.

## 5 Learn towards a low-temperature en-tropy state

Before exploiting a model’s self knowledge **p** to regularize the learning process, adjusting the knowledge’s entropy is vital. Deep model calibration [22] finds that modern neural networks are not well calibrated, and a model’s confidence can be aligned with its accuracy on a calibration dataset after training. A simple way is scaling a model’s output logits with a temperature before softmax normalisation [22, 30].

Without entropy adjustment, a deep neural network is significantly less confident when the training data has more severe noise and the confidence metrics are presented in Fig. 2 and Table 9. Therefore, we apply a low temperature *T* to decrease the entropy of self knowledge. Finally, other than using **p** to correct labels, we use

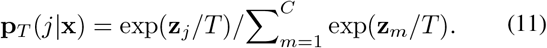

An annealed temperature (denoted by AT, 0 *< T <* 1) works best together with ProSelfLC, which trains a model towards a curated low-entropy target state.

For a comprehensive analysis, we also study AT integrated with other target modification approaches in the Section 6.5. AT boosts low-entropy (i.e., high-confidence) rewarding algorithms (i.e., Boot-soft and ProSelfLC) but does not help the low-entropy penalising method (i.e., CP).

## 6 Experiments

In deep learning, due to the stochastic batch-wise training scheme, small implementation differences (e.g., random accelerators like cudnn and different frameworks like Caffe [31], Tensorflow [1] and PyTorch [57]) may lead to a large gap of final performance. Therefore, to compare more properly, we re-implement CCE, LS and CP. Regarding Self LC methods, we re-implement Boot-soft [62], where *ϵ* is fixed throughout training. We do not re-implement stage-wise Self LC and KD methods, e.g., Joint Optimisation and Tf-KD_*self*_ respectively, because time-consuming tuning is required. In addition, our ProSelfLC can also be treated as an iteration-wise Self LC method. For an exact reproducibility and an entirely fair comparison, we fix the random seed and do not use any random accelerator in all experiments. By default, in clean and synthetic noisy cases, we train on 80% training data (corrupted in synthetic noisy cases) and use 20% trusted training data as a validation set to search all hyperparameters, e.g., Γ, *ϵ, B, T* and settings of an optimiser. Note that Γ and an optimiser’s settings are searched first and then fixed for all methods. Finally, we retrain a model on the entire training data (corrupted in synthetic noisy cases) and report its accuracy on the test data to fairly compare with prior results. In real-world label noise, the used datasets have a separate clean validation set. Here, a clean dataset is used only for validation, which is generally necessary for any method and differs from the methods [29, 32, 46, 63, 74, 75, 94, 107] that use a clean dataset to train a network’s learnable parameters.

### 6.1 Compare with baselines on clean CIFAR-100

#### Dataset and training details

CIFAR-100 [36] has 20 coarse categories and 5 fine classes in a coarse class. There are 500 and 100 images per class in the training and testing sets, respectively. The image size is 32 × 32. We apply simple data augmentation [28], i.e., we pad 4 pixels on every side of the image, and then randomly crop it with a size of 32 × 32. Finally, this crop is horizontally flipped with a probability of 0.5. We train the widely used ShuffleNetV2 [51] and ResNet-18 [28]. SGD is used with its settings as: (a) a learning rate of 0.2; (b) a momentum of 0.9; (c) the batch size is 128 and the number of training iterations is 39k, i.e., 100 epochs. We divide the learning rate by 10 at 20k and 30k iterations. The weight decay is 2*e*-3 for ResNet-18 while 1*e*-3 for ShuffleNetV2 because ResNet-18 has a larger fitting capacity.

#### Result analysis

We check the methods’ sensitivity to hyper-parameters, which generally makes more sense than that to random seeds. We report the mean and standard deviation of multiple hyper-parameters other than random seeds. Therefore, CCE has one run. While for LS, CP and Boot-soft, we run several different *ϵ* and *T*. Analogously, for ProSelfLC, we run several *B* and *T*. In detail, *ϵ* ∈ [0.125, 0.25, 0.375, 0.50], *T* ∈ [1.0, 0.8, 0.6, 0.4], *B* ∈ [20, 16, 12, 8]. The hyper-parameter space size is the same for each method except for CCE and LS. We report the mean and standard deviation of the top three results in Table 3. Compared with the baselines, ProSelfLC performs the best for both networks. First, we do not observe a big enough difference of those methods in the clean setting. Therefore, we focus on noisy scenarios hereafter. Second, the sensitivity to hyper-parameters is also small. Therefore, we do not report the standard deviation in the Table 4. Instead, we further discuss hyper-parameters in the Section 6.5.

**TABLE 3:**
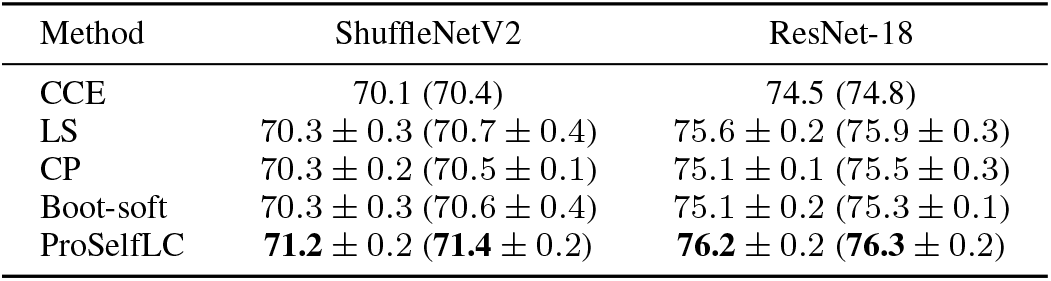
Test accuracy (%) on CIFAR-100 clean test set in the clean setting. We put the intermediately obtained best accuracy in the bracket. A lower drop from an intermediate best accuracy to the final one can be interpreted as a learner’s higher robustness against a long time being exposed to the training data. We bold the best results. The criteria of “best” is a highest intermediate accuracy followed with a highest final accuracy.

**TABLE 4:**
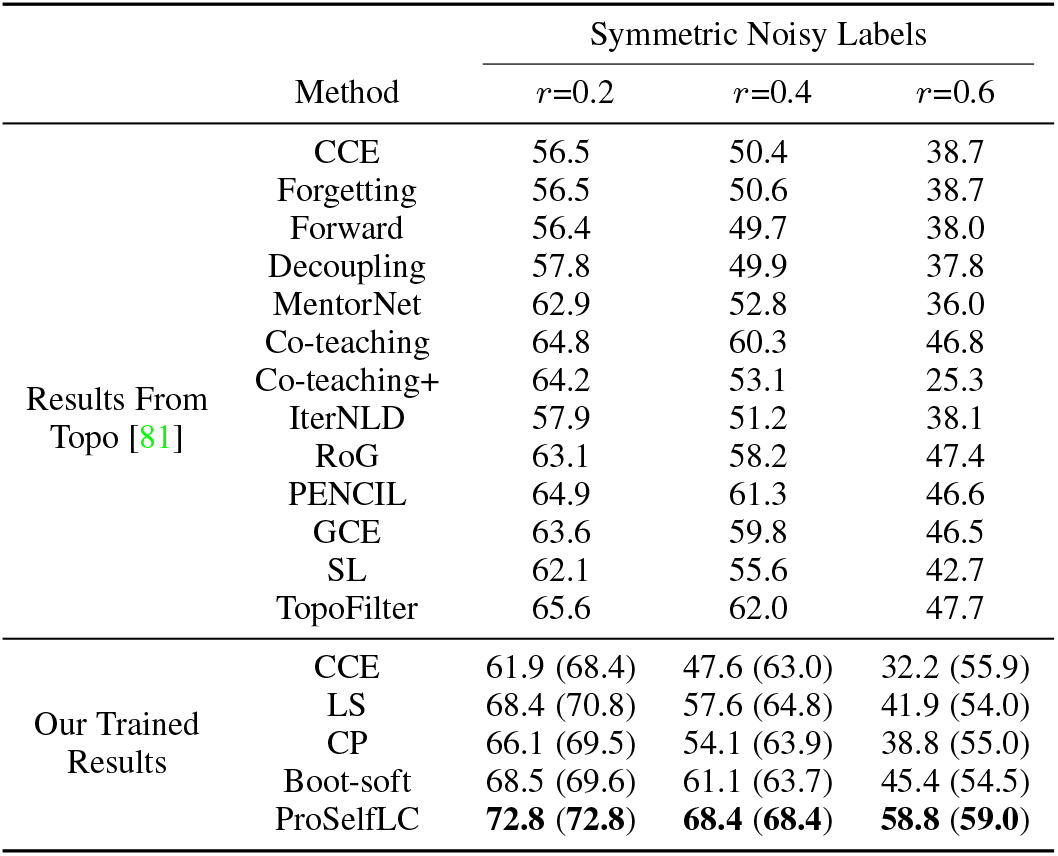
Test accuracy (%) on CIFAR-100 clean test set. The train set has a symmetric label noise rate of *r*. All compared methods use ResNet-18. We show the final accuracy and put the intermediate best one in corresponding bracket. A lower drop from the intermediate best to the final can be interpreted as a model’s higher robustness against a long time being exposed to the noise. We bold the best results.

### 6.2 Compare with the state-of-the-art methods on noisy CIFAR-100

#### Generating noisy train labels

(1) Symmetric label noise: the original label of an image is uniformly changed to one of the other classes with a probability of *r*. (2) Asymmetric label noise: we follow [79] to generate asymmetric label noise. Within each coarse class, we randomly select two fine classes *A* and *B*. Then we flip *r* × 100% labels of *A* to *B*, and *r* × 100% labels of *B* to *A*. We remark that the overall label noise rate is smaller than *r*.

#### Competitors^1^

We compare with the results reported recently in SL [79] and Topo [81]. Forward is a loss correction approach that uses a noise-transition matrix [58]. GCE denotes generalised cross entropy [106] and SL is symmetric cross entropy [79]. They are robust losses designed for solving label noise. Regarding the other robust loss functions including focal loss (FL) [48], NLNL [35], and normalised losses [52, 76, 77], according to the experimental report in active passive loss (APL) [52] where a deeper ResNet-34 is used though, their results are much worse than ours. Therefore, we do not compare with them in the table. Similarly, although TVD [103] uses ResNet-18, its reported results are much lower and not compared in the table. The other recent approaches are Forgetting [4], Decoupling [54], MentorNet [32], Co-teaching [25], Co-teaching+ [96], IterNLD [78], RoG [41], PENCIL [95] and TopoFilter [81]. Tf-KD_*reg*_ [97], SSKD [89] and Li’s LC [47] are three Self LC methods.

#### Results analysis

Training details are the same as Section 6.1. For all methods, we report their final results when training terminates. Therefore, we test the robustness of a model against not only label noise, but also a long time being exposed to the noise. In Table 4, we observe that: (1) ProSelfLC outperforms all the baselines, which is significant in most cases; (2) By default, we use AT and better baseline results are obtained. Despite that, our ProSelfLC further improves the standard Self LC (i.e., Boot-soft) and is the best of all. In addition, we visualize and comprehend the dynamic learning statistics of ProSelfLC versus baselines in Fig. 3, which clarifies why ProSelfLC works better. According to Table 5, ProSelfLC is superior to other recent Self LC methods.

**TABLE 5:**
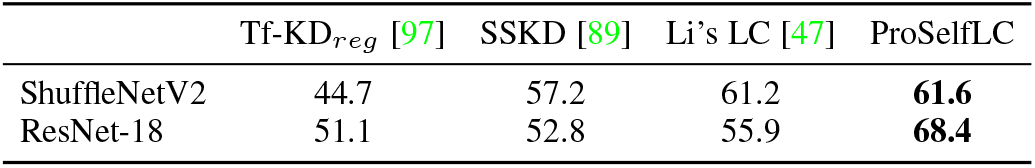
Test accuracy (%) on CIFAR-100 clean test set when the train set has 40% symmetric label noise. The Self LC (i.e., Self KD) baselines are implemented in [47] to test their robustness under label noise. The best results are bolded.

#### Revising the semantic class and similarity structure

In Fig. 3b and Fig. 3c, we show dynamic statistics of different approaches on fitting wrong labels and correcting them, respectively. ProSelfLC is much better than its counterparts. High semantic class correction means that the learned similarity structure revises the semantic class and similarity hierarchy corrupted by the noise.

#### To redefine and reward a low-temperature entropy state

On the one hand, LS and CP work well, being consistent with prior claims. In Fig. 3d and Fig. 3e, the entropies of both clean and noisy subsets are much higher in LS and CP, correspondingly their generalisation is the best except for ProSelfLC in Fig. 3f. On the other hand, ProSelfLC has the lowest entropy while performs the best, which proves that a learner’s confidence does not necessarily weaken its generalisation performance. Instead, a model needs to be cautious about what to be confident in. According to Fig. 3b and Fig. 3c, ProSelfLC has the lowest wrong fitting and highest semantic class correction, which indicates that the learned model reaches a low-entropy target state redefined by corrected labels.

### 6.3 Outperform the state-of-the-art methods on real-world noisy Clothing1M and Food-101N

#### Datasets

(1) *Clothing1M* [87] has about 1 million images of 14 fine-grained classes from shopping websites and around 38.46% label noise in the training data. Noisy labels are generated from description texts. The exact noise structure is unknown. In addition, Clothing1M is highly imbalanced, with images per label ranging from 18,976 to 88,588. The 14 classes are fine-grained and challenging to classify. A recent work Class2Simi [82] merges two similar classes into one class. To compare with the majority algorithms under the same setting, we do not merge classes. We train models only on the 1M noisy data and validate on clean test data. (2) *Food-101N* [42] has 101 fine-grained categories and contains 310K train images with noisy labels. The 25k curated test images are from the original Food-101 dataset [8]. In the train set, Food-101N [42] has 55k extra “verification labels” (55k VLs) which annotate whether noisy labels are correct or not. Following the recent papers [13, 26, 102, 105], we train on the more noisy dataset Food-101N [42] and validate on the original clean test data [8]. As a reference, although the recent robust early-learning method CDR [83] uses the less noisy Food-101 train data, its test accuracy is 86.36% and lower than ours.

#### Competitors

We compare with (1) sample selection methods including Search to Exploit (S2E) [93], CNLCU-S [84], TopoFilter [81], COnfidence REgularized Sample Sieve (CORES^2^) [14]; (2) Noise transition matrix estimation algorithms including Reweight [50], T-Revision [86], PTD-F-V [85], IF-F-V [33], kMEIDTM [13], TVD [103], VolMinNet [45], High-Order-Consensus (HOC) [109]; (3) Co-training methods, i.e., JoCoR [80] and Co-teaching [25]; (4) Meta-learning approaches, i.e., Meta-Cleaner [102] and Meta-Learner (MLNT) [44]; (5) Iterative label correction methods including Joint-Optim [73], P-correction [95] and PLC [105]; (6) Example weighting algorithms, i.e., MentorNet [32] and SIGUA [23]; (7) Compounded methods (linked by +): MD+DYR+SH [3] comprises three techniques, i.e., dynamic mixup (MD), dynamic bootstrapping together with label regularisation (DYR) and soft to hard (SH). MD+DYR+SH models sample loss with BMM [53] and applies Mixup [100]. Covariance-Assisted Learning (CAL) [108] exploits CORES^2^ [14] and Mixup. In the recent work [11], stochastic label noise (SLN) is proposed to regularize the training. Momentum (MO) denotes a model’s parameters are updated in a moving average way while LC means label correction. DivideMix [43] combines a set of strategies, including semi-supervised learning following dataset division and networks co-training. DivideMix used a different training schedule. Therefore, for a fair comparison, we compare with the DivideMix recently reproduced at SLN [11] from its released official implementation. ELR [49] denotes early-learning regularization. The recent ILFC [7] is not compared because it uses extra clean dataset and trains ResNet18. Nested co-teaching [12] trains ResNet18 and uses a very large batch size of 448. DAT [61] trained ResNet50 but used a batch size of 256. UniCon [34] integrates Auto-augment [15], contrastive learning and semi-supervised training, thus having slightly better results than us. The other methods have been introduced heretofore.

#### Experimental details

On both datasets, the ResNet-50 is pretrained on ImageNet and publicly available in PyTorch [57]. For Clothing1M, we follow the recent settings in [11, 105, 108] and use a small batch size of 32. The other training details are similar to Section 6.1 with small changes: we start with a learning rate of 0.01 and use a weight decay of 0.02. They are chosen according to the separate clean validation set. For Food-101N, we follow the same settings as the recent work [105]. The batch size is 128 and we train 72k iterations. We report the mean and standard deviation results of three random trials as in [11, 105].

#### Results analysis

In Table 6 of Clothing1M results, for both label-imbalanced data and label-balanced data, ProSelfLC has the highest accuracy, which demonstrates its effectiveness against real-world asymmetric and instance-dependent label noise. In Table 7, the results of Food-101N confirm again that ProSelfLC is superior to existing algorithms. We remark that Clothing1M and Food-101N contain fine-grained categories, thus being challenging. ProSelfLC obtains the state-of-the-art performance on both.

**TABLE 6:** Test accuracy (%) on the real-world noisy dataset Clothing1M, which contains asymmetric noise [95] and instance-dependent noise [7, 61]. We note that some approaches use the full training set while others sample a label-balanced subset. As two practices affect the performance a lot, we group the results into two columns for a clear and fair comparison. For the sampled noisy training data, each label has 18976 images, leading to about 260k images in total. * indicates online label-balanced sampling for each mini-batch. The first two blocks present the results from ICLR 2022 papers [84] and [33], respectively. The fourth block contains the results from [11]. Results of the third block are from multiple recent papers, as noted in the second column. When one method is reported in different papers and has different results, we keep the highest one only.

| Method | Where result was compared | Full training data: label-imbalanced | Sampled training data: label-balanced |
| --- | --- | --- | --- |
| S2E | [84] | 68.03 | – |
| MentorNet | [84] | 67.25 | – |
| SIGUA | [84] | 65.37 | – |
| JoCoR | [84] | 69.06 | – |
| CNLCU-S | [84] | 71.57 | – |
| Reweight | [33] | 70.82 | – |
| T-Revision | [33] | 71.27 | – |
| PTD-F-V | [33] | 70.62 | – |
| ELR | [33] | 71.86 | – |
| IF-F-V | [33] | 72.29 | – |
| MD+Dyr+SH | [3, 43, 81] | 71.00 | – |
| TVD | [103] | 71.65 | – |
| Meta-Cleaner | [43, 102] | 72.50* | – |
| Meta-Learner | [43, 44, 81, 105] | 73.47 | – |
| Joint-Optim | [43, 44, 49, 73, 81, 95, 102, 105] | 72.23 | – |
| kMEIDTM | [13] | 73.34 | – |
| P-correction | [43, 81, 95, 105] | – | 73.49 |
| PLC | [105] | – | 74.02 |
| TopoFilter | [81] | – | 74.10 |
| VolMinNet | [45] | – | 72.42* |
| HOC | [109] | – | 73.39* |
| CORES<sup>2</sup>+Mixup | [13, 14, 108, 109] | – | 73.24* |
| CORES<sup>2</sup>+Mixup+CAL | [13, 108] | – | 74.17 |
| CE | [11, 14, 33, 58, 81, 95, 102, 105, 108, 109] | 68.94 | 71.12 ± 0.32 |
| Forward | [11, 33, 44, 45, 58, 81, 95, 102, 105, 109] | 69.84 | 71.28 ± 0.27 |
| Backward | [11, 58] | 69.13 | 71.03 ± 0.33 |
| Co-teaching | [11, 109] | 70.19±0.28 | 72.14 ± 0.28 |
| DivideMix | [11] | – | 73.81±0.41 |
| SLN | [11] | 70.42 ± 0.34 | 72.95 ± 0.31 |
| SLN+MO | [11] | 71.15 ± 0.21 | 72.98 ± 0.15 |
| SLN+MO+LC | [11] | 72.61 ± 0.23 | 74.08 ± 0.18 |
| ProSelfLC | Ours | **73.93 ± 0.02** | **74.48 ± 0.01** |

**TABLE 7:** Test accuracy (%) on the real-world noisy dataset Food-101N, composed of 101 fine-grained food classes.

| Method | Where result was compared | Train Data | Accuracy |
| --- | --- | --- | --- |
| CCE | [13, 42, 102] | Food-101N | 81.44 |
| CleanNet ($w_{hard}$) | [13, 26, 42, 102] | Food-101N+55k VLs | 83.47 |
| CleanNet ($w_{soft}$) | [13, 26, 42, 102, 105] | Food-101N+55k VLs | 83.95 |
| Meta-Cleaner | [102] | Food-101N | 85.05 |
| Multi-Prototypes | [13, 26] | Food-101N | 85.11 |
| PLC | [105] | Food-101N | 85.28±0.04 |
| DivideMix | [13] | Food-101N | 84.39 |
| kMEIDTM+DivideMix | [13] | Food-101N | 85.61 |
| ProSelfLC | Ours | Food-101N | **86.62±0.01** |

### 6.4 Train robust transformers on noisy protein classification datasets

We follow the recent ProtTrans to do experiments on protein classification [17]. Training deep models to predict a protein’s properties is challenging as the length of amino acid sequences varies from several tens to multiple thousands [2, 17, 64]. Some recent approaches crop amino acid sequences to decrease training time and GPU memory consumption [2, 17, 64]. In this work, we find that cropping input sequences adds noise to model training as some proteins have important functional regions interspersed across the protein length [2]. We empirically demonstrate this by training models on the cropped proteins. In addition to cropping noise, we further design high-noise experiments by including unlabelled proteins. Conceptually, it is semi-supervised learning. We bridge semi-supervised learning and label-noise learning by assigning random labels to those unlabelled proteins, so that we establish the synthetic dataset to validate ProSelfLC for training robust protein transformers against label noise.

#### Datasets

The DeepLoc train set used in [17] contains 6,622 proteins whose are annotated to be membrane (i.e., they were found on the membrane), water-soluble (i.e., they were from the lumen of the organelle), or unknown (missing information about where they were found) [2]. Specifically, there are 1,518 transmembrane proteins, 2,227 water-soluble proteins and 2,877 proteins with unknown labels. There are 1,842 proteins in total and 1,087 proteins with known labels in the test set. We present the cropping noise and synthetic symmetric label noise as follows:

- *Cropping noise*. We train on proteins with known labels. Their length ranges from 40 to 13,100, with a median of 434. Therefore, we truncate proteins longer than 434 at the end so that all amino acid sequences have a length no longer than 434. The goal of this setting is to validate whether ProSelfLC could be robust to cropping noise if there is a practical need to crop sequences for speeding up training and reducing GPU memory requirement.
- *Cropping noise+Label noise*: First, we keep the cropping noise as we crop proteins to decrease training time and reduce GPU memory requirement. Second, we add symmetric label noise by assigning uniform random labels to 2,877 unlabelled proteins. Cropping noise together with label noise makes the noise level high. The objective is to evaluate whether ProSelfLC could be robust to severe noise when it is expensive to remove it in practice.

#### Network and training details

We train sequence transformers to classify a single amino acid sequence (without using homology information at all) to be either transmembrane or water-soluble. The transformer network is a subnet of ProtBert-BFD [17], a protein language model pretrained on BFD-100 dataset [69, 70]. According to [17], to use a larger batch size, ProtBert-BFD was first trained on sequences with maximum length of 512, then tuned on sequences with maximum length of 2k. We name this subnet ProtBert-H16-D6, where the D6 denotes its depth is 6 (D6), i.e., a stack of 6 hidden transformer layers. H16 means that in each transformer layer, the number of transformer blocks (a.k.a., heads) is 16. The released model has a depth of 30, so that our subnet is 5 times shallower. We choose to train this subnet mainly because it requires a small-memory GPU and is faster to train. In addition, it benefits little from pretraining, so it indicates our algorithm can be applied to train sequence transformers from scratch. Finally, we will release this subnet and make it convenient to reproduce our results with a 16GB GPU machine. We use a batch size of 32 and the SGD optimiser. The weight decay is 0.0001. For cropping noise, a starting learning rate of 0.02 is used. When noise rate is high, i.e., cropping noise+label noise, we use a smaller starting learning rate of 0.01. We train 40 epochs in total. We stress that the generic hyperparameters are coarsely searched by visualizing the statistical training curves, i.e., without brute-force and confidently fitting all training data as noise exists. Besides, we report the metrics of the final model when training terminates other than select the best intermediate model, leaving our reported metrics less biased.

#### Results analysis and discussion

We present the accuracy and confidence metrics of ProSelfLC and baseline algorithms in Table 8. ProSelfLC’s fitting of the noisy train set is much lower than that of CCE, which indicates that ProSelf does not overfit the training set. This observation is obvious for both noise types. Furthermore, ProSelfLC learns and generalises better and more confidently compared with other widely used baselines. Our experiments demonstrate that the misleading effect by cropping noise can be alleviated by ProSelfLC, as ProSelfLC’s performance (92.4%) is even slightly better than the DeepLoc ensemble model (92.3%) [2] and the large transformer model (91.0%) [17]. With the high label noise added, though 2,877 out of 6,622 proteins have random labels, the generalisation performance of ProSelfLC decreases little from 92.4% to 92.2%, which confirms that ProSelfLC can be a robust solution when it is expensive to remove severe noise in practice.

**TABLE 8:** Results of robust protein classification against cropping noise and symmetric label noise. For comprehensiveness, we report accuracy and confidence metrics on both train and test sets. We show two confidence metrics: 1− *H*(**p**)*/H*(**u**) and max_*j*_ **p**(*j* | **x**). We use the final model when training ends without selecting the intermediate best ones. The highest accuracy or confidence in each row is bolded.

| Noise type | Metric | Data | CCE | LS | CP | Boot-soft | ProSelfLC |
| --- | --- | --- | --- | --- | --- | --- | --- |
| Cropping noise | Accuracy | Test | 88.7 | 90.4 | 90.9 | 90.4 | **92.4** |
|  |  | Train | **99.3** | **99.3** | 98.8 | 95.9 | 91.8 |
| | $1 - \frac{H(\mathbf{p})}{H(\mathbf{u})}$ | Test | 86.9 | 59.5 | 83.8 | 89.0 | **96.4** |
|  |  | Train | 89.0 | 63.3 | 82.0 | 92.6 | **97.6** |
| | $\max_j \mathbf{p}(j\vert \mathbf{x})$ | Test | 96.4 | 90.1 | 95.5 | 96.8 | **99.2** |
|  |  | Train | 97.6 | 92.2 | 95.7 | 98.5 | **99.7** |
| Cropping noise + Label noise | Accuracy | Test | 84.2 | 89.4 | 89.4 | 89.6 | **92.2** |
|  |  | Train | **84.8** | 75.2 | 72.8 | 77.2 | 75.7 |
| | $1 - \frac{H(\mathbf{p})}{H(\mathbf{u})}$ | Test | 65.3 | 38.7 | 37.4 | 57.6 | **96.1** |
|  |  | Train | 42.0 | 20.0 | 18.0 | 31.3 | **95.2** |
| | $\max_j \mathbf{p}(j\vert \mathbf{x})$ | Test | 88.9 | 80.4 | 78.4 | 86.0 | **99.2** |
|  |  | Train | 80.7 | 70.0 | 67.5 | 74.9 | **99.1** |

### 6.5 Ablation studies

#### Normal-temperature entropy state versus low-temperature entropy state

For CP, Boot-soft and our ProSelfLC, the target state’s entropy can be adjusted by the temperature. We denote annealed temperature by AT. We study the normal-temperature state (i.e., without AT) versus the low-temperature state (i.e., with AT) and display their results in Table 9. Generic training parameters are the same for all methods. We observe: (1) Compared with the baseline CCE, the confidence-penalty approaches (LS and CP) indeed learn better and lower-confidence models, which is consistent with the proposed motivations of them; (2) However, confidence-reward algorithms (Boot-soft and ProSelfLC) can perform better. ProSelfLC with AT generalises the best with the highest confidence, i.e., the lowest entropy; (3) Across networks and noise rates, CP is less sensitive to target-state entropy while Boot-soft is the most sensitive. This demonstrates that target-state entropy is also very important for standard label correction (i.e., Boot-soft). For ProSelfLC, AT improves the performance consistently and reaches a curated low-temperature entropy state. Therefore, for both Boot-soft and ProSelfLC, we use AT by default in other experiments.

**TABLE 9:**
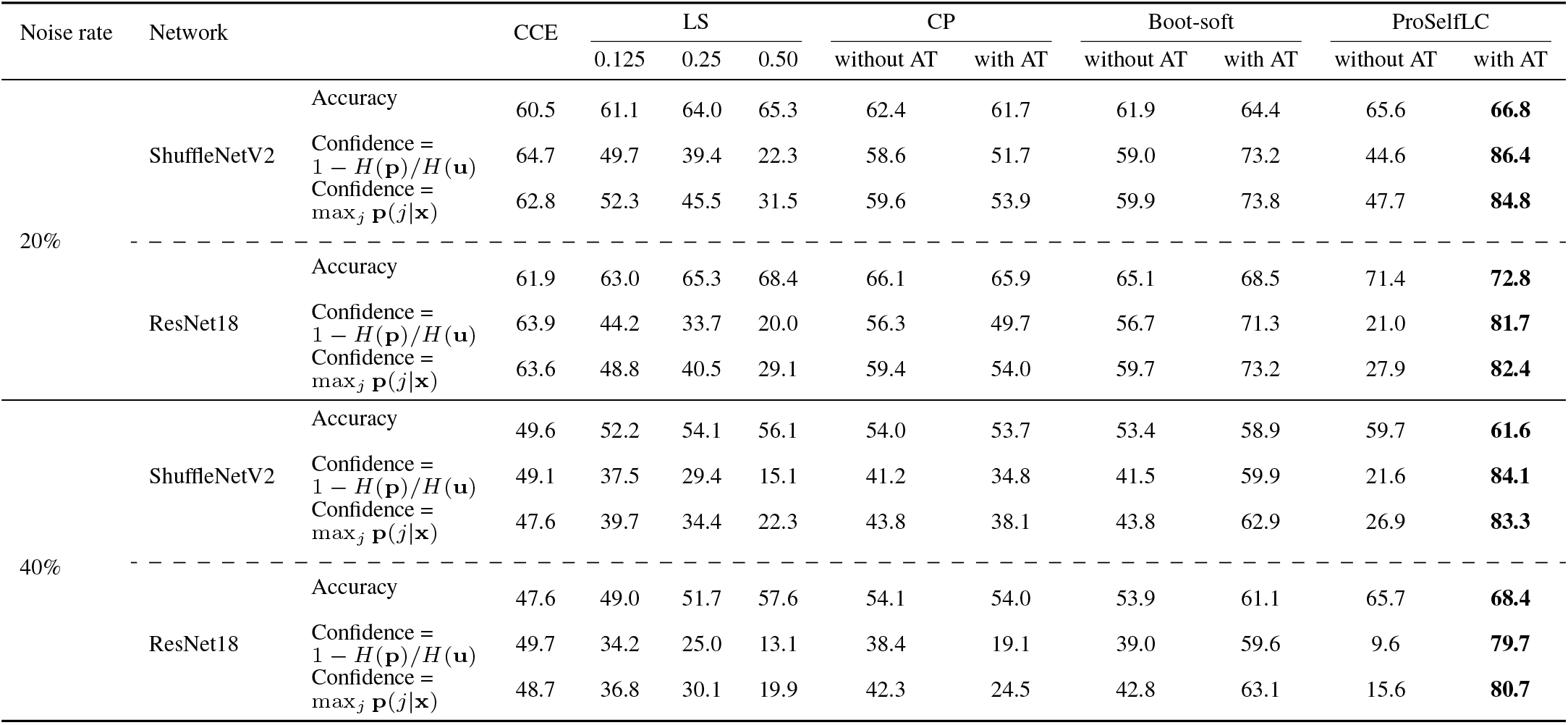
Results of target modification methods without/with annealed temperature (AT) for curating the target-state entropy. We train models on CIFAR-100 whose training labels contain symmetric noise. We do not select the intermediate best models and report the generalisation accuracy and confidence metrics on the clean test set when training terminates. There are two confidence measurements: 1 − *H*(**p**)*/H*(**u**) and max_*j*_ **p**(*j*|**x**). The highest accuracy or confidence of each row is bolded.

**TABLE 10:**
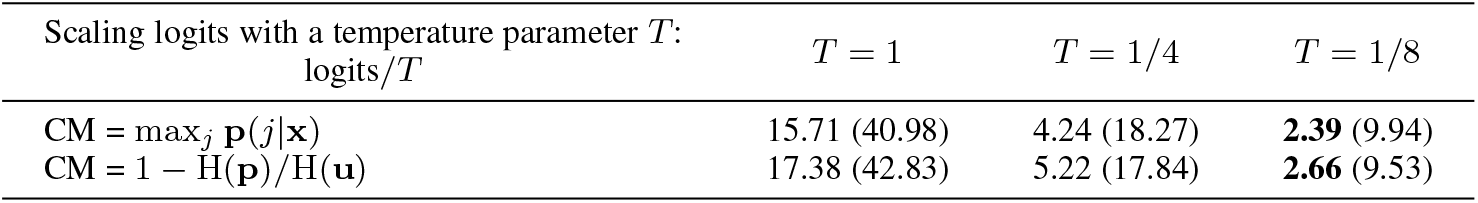
ECE results of multiple combinations of logits scaling (logits*/T*) and confidence metrics (probability and entropy).

| Scaling logits with a temperature parameter $T$:<br>logits/$T$ | $T = 1$ | $T = 1/4$ | $T = 1/8$ |
| --- | --- | --- | --- |
| CM = $\max_j \mathbf{p}(j\vert \mathbf{x})$ | 15.71 (40.98) | 4.24 (18.27) | **2.39** (9.94) |
| CM = $1 - H(\mathbf{p})/H(\mathbf{u})$ | 17.38 (42.83) | 5.22 (17.84) | **2.66** (9.53) |

#### Hyper-parameters space of *B* and *T*

To be more comprehensive, we do experiments on CIFAR-100 without/with symmetric lable noise. Additionally, we train both ShuffleNetV2 and ResNet18. In those experiments, we only change *B* and *T*. All other training parameters are the same. We use the model when training ends without selecting the intermediate ones. According to the Fig. 4, on both networks and both noise rates, generally, the performance is more sensitive to *T* when *B* is large. This confirms a human’s intuitive concept that if we trust a learner itself in a faster speed, the confidence adjustment of this learner’s predictions becomes more crucial. When the noise rate increases, better results can be obtained by using a relatively smaller *T* to optimise the model towards a low-temperature entropy state.

**Fig. 4:**
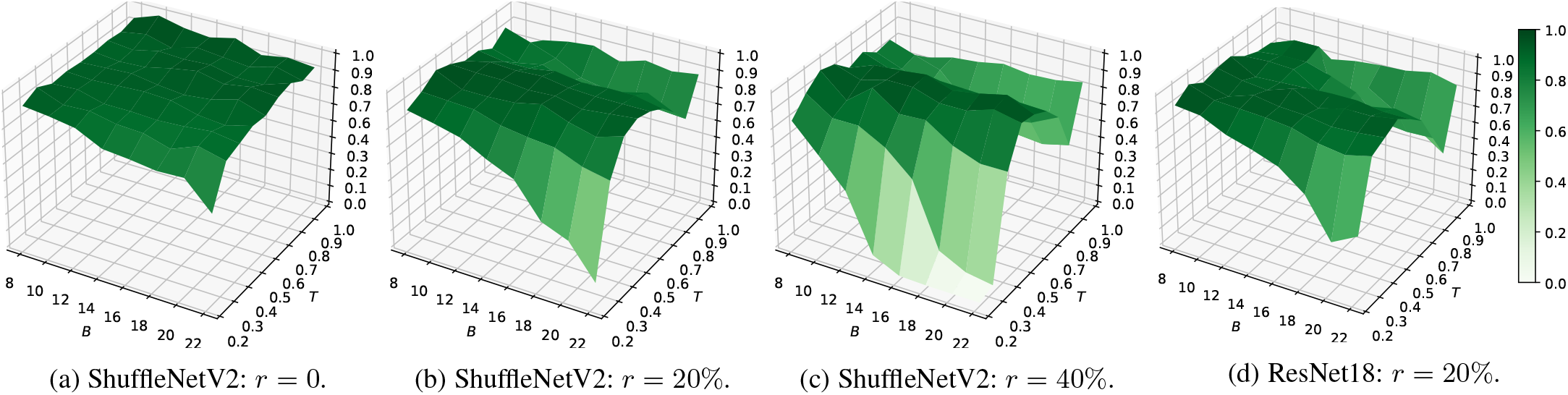
Grid analysis on the hyper-parameters space of *B* and *T*. We use CIFAR-100 with a symmetric label noise rate of *r*. For a clearer visualization, in each subfigure, all results are plotted in an exponential ratio to the best one.

#### Self trust schemes

We study the differences of four self trust schemes described in Section 4. When *ϵ* _ProSelfLC_ is constant at training, it degrades to Boot-soft. According to the Fig. 5, we observe that (1) Compared with “constant”, *g*(*t*) outperforms Boot-soft in three metrics except for sacrificing fitting clean train subset a lot. (2) Compared with “constant” and *g*(*t*), *g*(*t*) ∗ (1− H(**p**)*/*H(**u**)) and *g*(*t*) ∗ max_*j*_ **p**(*j* |**x**) are better in balancing fitting and generalisation. By default, in all other experiments, we use *g*(*t*) ∗ (1− H(**p**)*/*H(**u**)) due to its slightly better results.

**Fig. 5:**
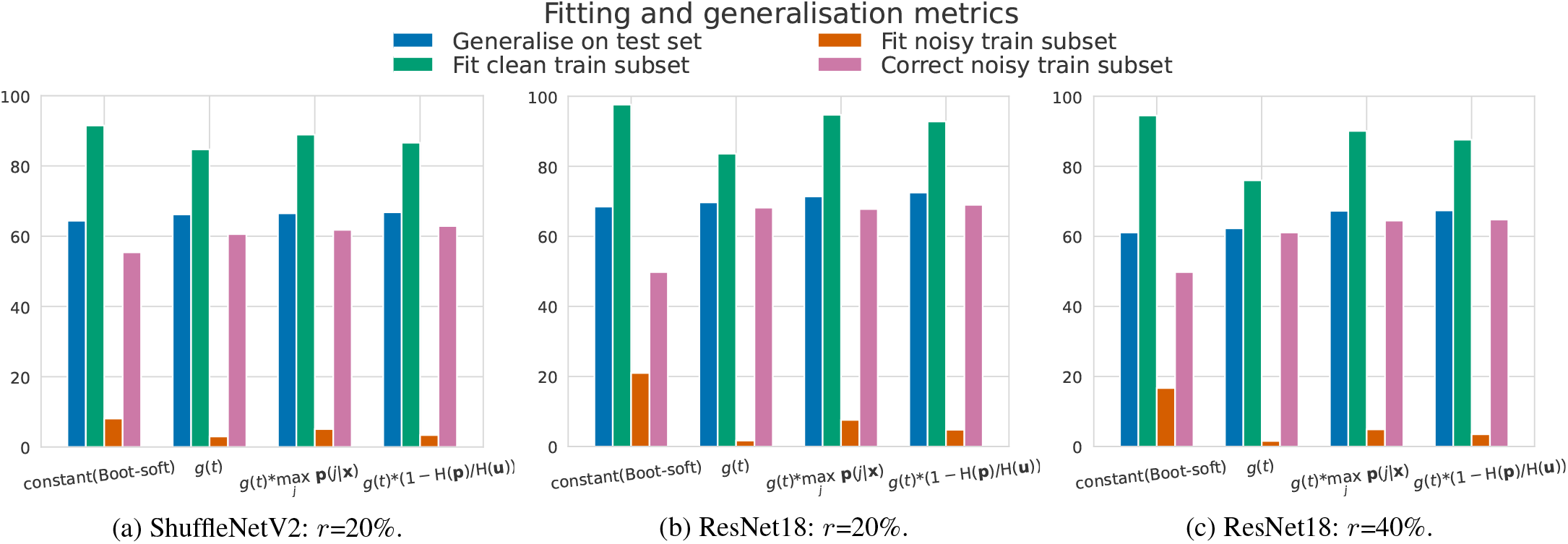
Results of computing *ϵ* _ProSelfLC_ by four self trust schemes, denoted in the horizontal axis. Experiments are done on CIFAR-100 with symmetric label noise. A subfigure’s caption describes the used network and noise rate *r*. We report four metrics (%) and one coloured bar per metric along the vertical axis. *For fitting noisy train subset, a lower value is better. For other three metrics, a higher value is better*. According to the finding of AT’s effectiveness for boosting Boot-soft and ProSelfLC in the Table 9, *we add AT on top of every self trust scheme*. Therefore, we observe all schemes have competitive results and their performance gaps become smaller. At training, a learner is not given whether a label is noisy or not. We use the final model when training ends.

We further discuss post-training model calibration [22] in Appendix B, and the changes of entropy and *ϵ* _ProSelfLC_ during training in Appendix C.

## 7 Conclusion

We present a thorough mathematical study on several target modification techniques. Through analysis of entropy and KL divergence, we reveal their relationships and limitations. To improve and endorse self label correction, we propose ProSelfLC. Extensive experiments prove its superiority over existing methods under clean and noisy settings. ProSelfLC enhances the similarity structure information over classes, and rectifies the semantic classes of noisy label distributions.

Inspired by the finding that a deep model is of high entropy under severe noise, ProSelfLC is the first approach to trust low-temperature self knowledge progressively and adaptively. Pro-SelfLC redirects and promotes entropy minimisation, which is in marked contrast to recent practices of confidence penalty [16, 59, 72].

Furthermore, this work presents a new insightful finding to complement a previous one “deep neural networks easily fit random labels [99]”: *Deep models fit and generalise significantly less confident when more random labels exist*.

**Xinshao Wang** is a senior researcher of Zenith Ai and a visit scholar of University of Oxford. He was a postdoctoral researcher at the Department of Engineering Science, University of Oxford after finishing his PhD at the Queens University of Belfast, UK. Xinshao Wang is working on core deep learning techniques with applications to visual recognition, disease prediction based on electronic health records, and protein engineering. Concretely, he has been working on the following topics: (1) Deep metric learning: to learn discriminative and robust representations for downstream tasks, e.g., object retrieval and clustering; (2) Robust deep learning: robust learning and inference under adverse conditions, e.g., noisy labels, missing labels (semi-supervised learning), out-of-distribution training examples, sample imbalance, etc; (3) Computer vision: video/set-based person re-identification; image/video classification/retrieval/clustering; (4) AI healthcare: electrocardiogram classification; (5) ML-assisted gene and protein engineering.

**Yang Hua** is presently a lecturer at the Queen’s University of Belfast, UK. He received his Ph.D. degree from Université Grenoble Alpes / Inria Grenoble Rhne-Alpes, France, funded by Microsoft Research Inria Joint Center. He won PASCAL Visual Object Classes (VOC) Challenge Classification Competition in 2010, 2011 and 2012, respectively and the Thermal Imagery Visual Object Tracking (VOTTIR) Competition in 2015. His research interests include machine learning methods for image and video understanding. He holds three US patents and one China patent.

**Elyor Kodirov** is a Lead Researcher at Zenith Ai and leads the PROEML team. He received his Ph.D. degree from Queen Mary University of London, UK. He was supervised by Tao Xiang. His research interests include machine learning, computer vision and NLP. Previously, he worked on computer vision with application to person reidentification/detection and face recognition. At present, his focuses on AI for bioinformatics and cheminformatics. He is working on challenging problems such as genetic expansion, protein engineering and molecular/protein representaion learning.

**Sankha Subhra Mukherjee** is a hands-on research leader who thinks deeply about the hardest problems in machine learning and delivers results on which innovative businesses have been created. Dr Mukherjee is a deep learning expert. His doctoral research at Heriot-Watt developed new deep neural network techniques which led to the cofounding of a high growth tech start-up, landmark publications, and patents. He had been the founding EVP of Research and later CSO for a high growth start-up leading breakthroughs in machine learning, recruiting and supervising a team of 20 world-class researchers. In his current role he is a co-founder and Chief Scientific Officer, leading a team of world class researchers to deliver breakthroughs in ML and AI driven cell engineering and synthetic biology.

**David A. Clifton** is a Professor of Clinical Machine Learning and leads the Computational Health Informatics (CHI) Lab. He is Official Fellow in AI & ML at Reuben College, a Research Fellow of the Royal Academy of Engineering, and a Fellow of Fudan University, China. He studied Information Engineering at Oxford’s Department of Engineering Science, supervised by Professor Lionel Tarassenko CBE. His research focuses on ‘AI for healthcare’. In 2018, the CHI Lab opened its second site, in Suzhou (China), with support from the Chinese government. In 2019, the Wellcome Trust’s first “Flagship Centre” was announced, which joins CHI Lab to the Oxford University Clinical Research Unit in Vietnam, focused on AI for healthcare in resource-constrained settings. He is a Grand Challenge awardee from the UK Engineering and Physical Sciences Research Council, which is an EPSRC Fellowship that provides long-term strategic support for nine “future leaders in healthcare.” He was joint winner of the inaugural “Vice-Chancellor’s Innovation Prize”, which identifies the best interdisciplinary research across the entirety of the University of Oxford.

**Neil M. Robertson** is CEO of Zenith Ai, and a Professor of Electrical Engineering and Computer Science at Queen’s University Belfast. He leads research into underpinning machine learning methods applied to diverse areas including synthetic biology, visual analytics and robotics. He started his career in the UK Scientific Civil Service (2000-2007), was the 1851 Royal Commission Fellow at Oxford University (2003–2006) in the Robotics Research Group and an academic at Edinburgh (2007-2016). His ML/AI research is extensive including UK major research programmes and a national doctoral training centre. He is the co/founder of three successful AI companies.

## Appendix A

### Proof of propositions

#### Proposition 3.

*Compared with CCE, LS and CP penalise entropy minimisation while LC reward it. Proof*. We can rewrite CCE, LS, CP, and LC from the viewpoint of KL divergence:

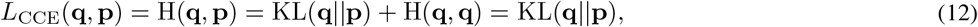

where we have H(**q, q**) = 0 because **q** is a one-hot distribution.

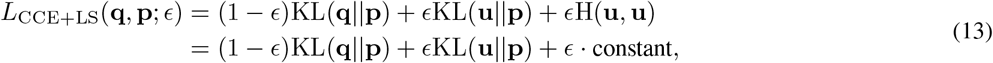

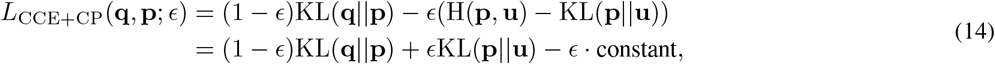

where H(**p, u**) = H(**u, u**) = constant. Analogously, LC in Eq (5) can also be rewritten:

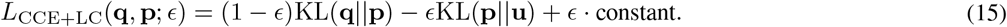

In LS and CP, both +KL(**u** || **p**) and +KL(**p** || **u**) pulls **p** towards **u**. While in LC, the term KL(**p** || **u**) pushes **p** away from **u**. □

#### Proposition 4.

*Proof*. In LS, the target is 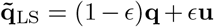. For any 0 ≤ *ϵ <* 1, the semantic class is not changed, because 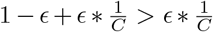. In addition, 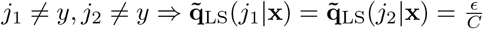.

In CP,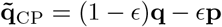. In terms of label definition, *CP is against intuition because these zero-value positions in* **q** *are filled with negative values in* 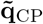. A probability has to be not smaller than zero. So we rephrase 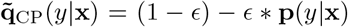, and ∀*j ≠ y*, 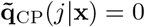by replacing negative values with zeros, as illustrated in Fig. 1a. □

## Appendix B

### Discussions on wrongly confident predictions and model calibration

1. *It is likely that some highly confident predictions are wrong. Will ProSelfLC suffer from an amplification of those errors?* First of all, ProSelfLC alleviates this issue a lot and makes a model confident in correct predictions, according to Fig. 3e together with 3b and 3c. ***Fig. 3e shows the confidence of predictions, whose majority are correct according to Fig. 3b and 3c***. In Fig. 3b, ProSelfLC fits noisy labels least, i.e., around 12% so that the correction rate of noisy labels is about 88% in Fig. 3c. Nonetheless, ProSelfLC is non-perfect. A few noisy labels are memorised with high confidence.
2. *How about the results of model calibration using a computational evaluation metric: Expected Calibration Error (ECE) [22, 56]?* Following the practice of [22], on the CIFAR-100 test set, we report the ECE (%, #bins=10) of ProSelfLC versus CCE, as a complement of Fig. 3. For a comparison, CCE’s results are shown in corresponding brackets. We try several confidence metrics (CMs), including probability, entropy, and their temperature-scaled variants using a parameter *T*. Though the ECE metric is sensitive to CM and *T*, ProSelfLC’s ECEs are smaller than CCE’s.

## Appendix C

### The changes of entropy statistics and *E*ProSelfLC at training

In Fig. 6, we visualise how the entropies of noisy and clean subsets change at training.

**Fig. 6:**
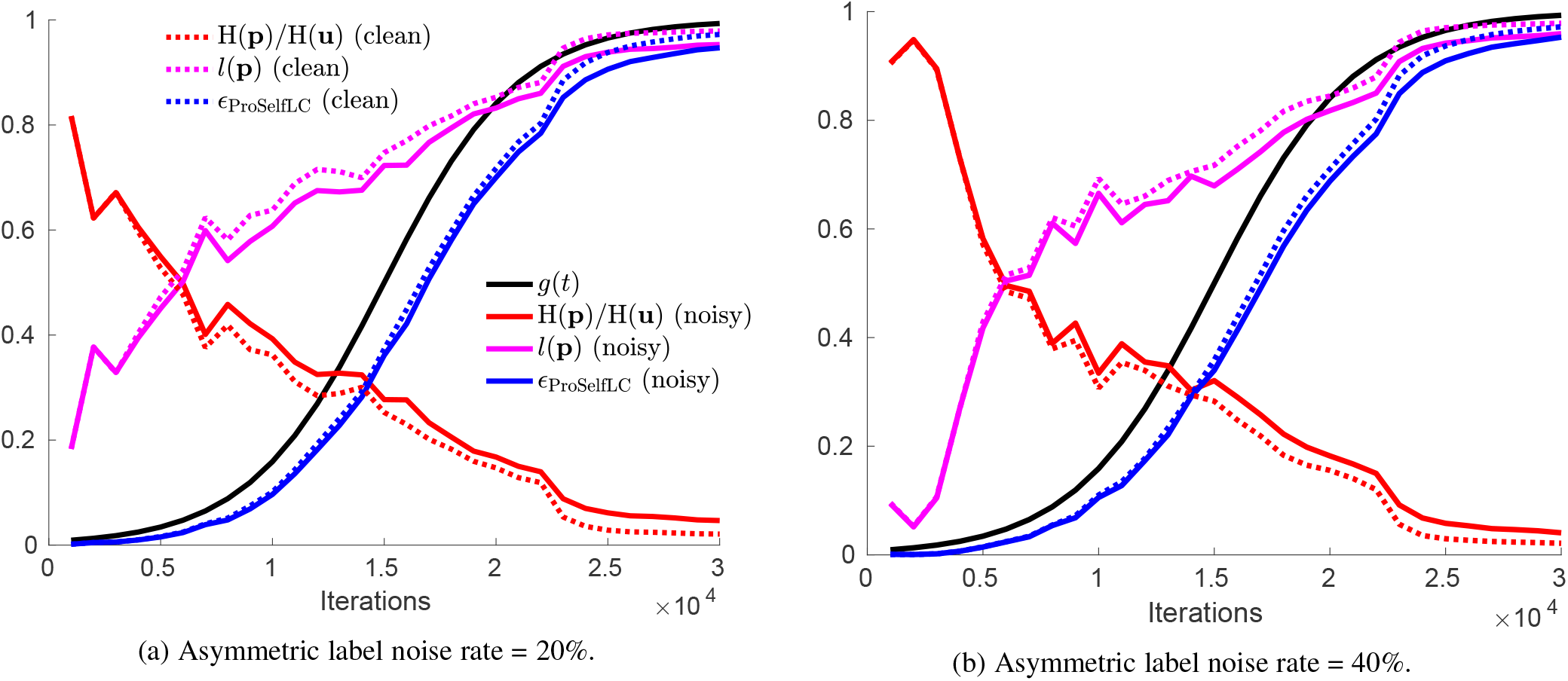
The changes of entropy statistics and *ϵ* _ProSelfLC_ at training. We store a model every 1000 iterations to monitor the learning process. For data-dependent metrics, after training, we split the corrupted training data into clean and noisy subsets according to the information about how the training data is corrupted before training. Finally, we report the mean results of each subset.

## Footnotes

1 We do not consider DisturbLabel [88], which flips labels randomly and is counter-intuitive. It weakens the generalisation because generally the accuracy drops as the uniform label noise increases.

